# Disruptions in glucose and amyloid-beta transport in mouse models manifesting metabolic syndrome

**DOI:** 10.64898/2026.08.15.741912

**Authors:** Lushan Wang, Geoffry L. Curran, Chaitanya Chakravarthi Gali, Andrew L. Zhou, Paul H. Min, Val J. Lowe, Karunya K. Kandimalla

## Abstract

Studies in humans and murine models have pointed towards a possible link between metabolic syndrome, which shows insulin resistance and metabolic dysregulation, and Alzheimer’s disease (AD) pathology marked by amyloid-beta (Aβ) accumulation at the BBB and hypometabolism in the brain. Yet, the underlying biological mechanisms by which metabolic syndrome affects these pathological changes in AD brain remain unknown. We hypothesized that insulin resistance is responsible for alterations in blood-brain barrier (BBB) transport of Aβ peptides and glucose. This hypothesis was tested by employing radiolabeled tracers (^125^I-Aβ40, ^125^I-Aβ42, and ^18^F-FDG) in high-fat diet (HFD)-fed mouse models that manifest metabolic syndrome. Further, we assessed alterations in the expression of various molecular mediators within the brain microcapillaries harvested from both low-fat diet (LFD)-fed and HFD-fed mice. As expected, our studies show that HFD-fed mice developed peripheral insulin resistance and obesity. In addition, HFD-fed mice demonstrated an increase in the influx rate of Aβ peptides and a reduction in ^18^F-FDG (a glucose surrogate) influx rate at the BBB compared to LFD-fed mice. These transport changes are associated with increase in the BBB endothelial expression of RAGE (receptor to traffic Aβ from plasma-to-brain) and a reduction of GLUT1 (glucose transporter) expression in HFD-fed mice compared to LFD-fed mice. Moreover, disruption in insulin signaling, as indicated by reduced pAKT and pERK expression, was observed in HFD-fed mice. Inhibiting AKT or ERK phosphorylation with specific inhibitors resulted in similar changes in Aβ and glucose uptake in polarized BBB endothelial cell monolayers in vitro. These results indicate that HFD induced metabolic syndrome may lead to BBB dysfunction, characterized by increased plasma-to-brain Aβ trafficking and diminished glucose transport at the BBB, thereby aggravating the expression of AD pathological hallmarks.

## INTRODUCTION

Metabolic syndrome, which manifests obesity, hyperinsulinemia, and dyslipidemia, is a major risk factor for late-onset Alzheimer’s disease (LOAD)^1^. Studies conducted in Alzheimer’s disease (AD) transgenic mouse models have demonstrated that metabolic syndrome induced by a high-fat diet (HFD) contributes directly to increased amyloid beta (Aβ) accumulation in peripheral organs and the brain of various AD transgenic mouse models, including PDGF-APP_Sw,Ind_^2^, APP_swe_/PS1_ΔE9_^3–5^, Tg2576^6–8^ and 3XTg-AD^9–12^ mouse models. In addition, an increase in the brain Aβ levels was observed in cross-mated AD and type-2 diabetes mellitus (T2DM) mouse models^13–16^.

The blood-brain barrier (BBB), a selective transport portal for molecules in and out of the brain, plays a major role in transporting glucose from plasma to the brain and in Aβ trafficking between the brain and plasma^17–19^. The receptor for advanced glycation end products (RAGE), located on the luminal side of the BBB, and low-density lipoprotein receptor-related protein 1 (LRP1), located on the abluminal side of the BBB, regulate Aβ transport in and out of the brain, respectively^19–22^. Furthermore, P-glycoprotein (P-gp), a member of the ATP-binding cassette (ABC) transporter family expressed on the luminal side of the BBB, is also implicated in facilitating brain Aβ clearance^23–25^.

Metabolic syndrome is shown to be associated with reduced cerebral metabolic rate of glucose (CMRglc) based on the studies conducted in animal models and in a limited number of human studies^26, 27^. Often, this reduction is linked to a decrease in the expression of glucose transporter 1 (GLUT1), the primary transporter of glucose at the BBB, as well as GLUT3, the primary glucose transporter in neurons^28–30^. Researchers have hypothesized that in metabolic syndrome, the peripheral tissues are exposed to higher glucose levels, but the brain glucose levels may be reduced due to the downregulation of GLUTs at the BBB^31, 32^. Although it seems to be a protective mechanism that reduces glucose exposure to the cerebrovascular endothelium, it may cause a severe deficiency in glucose availability to the brain.

To investigate the impact of metabolic syndrome on dysregulation of Aβ trafficking and glucose transport disruptions at the BBB, we examined the plasma distribution and brain influx kinetics of radiolabeled amyloid-beta (^125^I-Aβ40 or ^125^I-Aβ42) and ^18^fluorodeoxyglucose (^18^F-FDG) in mice maintained on low-fat diet (LFD) versus HFD. We further elucidated molecular changes associated with BBB trafficking disruptions in these mouse models.

## MATERIALS AND METHODS

### Materials

Aβ40 and Aβ42 peptides were procured from AAPPtec, LLC (Louisville, KY). The ^125^I-Na was obtained from PerkinElmer Life and Analytical Sciences (Boston, MA). Insulin (Novolin® or Humulin®) was purchased from Eli Lily (Indianapolis, IN).

### Animals

Female B6SJLF1/J mice (MMRRC Strain # 100012), aged two months, were acquired from the Jackson Laboratory (Bar Harbor, ME) and subsequently housed at the Mayo Clinic’s animal care facility. The mice were maintained in an environment with regulated humidity and temperature, unimpeded access to food and water, and an established 12-hour light-dark cycle. At the age range of 7-11 months, the mice were assigned to diets differing in fat content: either a high-fat diet (HFD) constituting 60% of calories from fat (D12492, Research Diets, Inc., New Brunswick, USA) or a low-fat diet (LFD) with 10% of calories from fat (D12450B, Research Diets, Inc., New Brunswick, USA). This dietary regimen was maintained for a duration of 13-16 weeks, during which body weights were systematically logged on a weekly basis. Insulin Tolerance Tests (ITT) were performed both prior to and after this feeding period. Before the experiment, the animals underwent overnight fasting for 12 hours. All experimental procedures adhered to the National Institutes of Health guidelines for laboratory animal care and received approval from the Mayo Clinic Institutional Animal Care and Use Committee (Mayo IACUC # A00002082-16-R22). Additionally, prior to the experiments, the mice were assessed to ensure they were in the diestrus phase of the estrous cycle ^33, 34^ to mitigate potential hormonal fluctuation impacts. The reported experiments were performed in compliance with the guidance for the Care and Use of Laboratory Animals outlined by the National Institute of Health^40^ as well as the ARRIVE guidelines 2.0 (Animal Research: Reporting in Vivo Experiments)^41^.

### ITT

Prior to the test, mice underwent a fasting period of 4 hours, after which they received an intraperitoneal injection of 0.2 milli-Units per gram of Humulin R insulin (Eli Lilly and Company, Indianapolis, IN) as described previously ^35^. Blood glucose levels of the mice were monitored at 0, 15, 25, 45, 60, and 90 minutes following the Humulin® injection and measured using the Precision Xtra Blood Glucose Monitoring System (Abbott Laboratories, Chicago, IL).

### Radioiodination of Aβ peptides and FDG

Aβ peptides were labeled with ^125^I radionuclide (PerkinElmer Life and Analytical Sciences, Boston, MA) using the chloramine-T procedure as described previously ^36, 37^. After dialysis against 0.01 M PBS at pH 7.4 (SigmaAldrich, St. Louis, MO) to remove unconjugated radionuclide, the purity of ^125^I-labeled Aβ was determined by trichloroacetic acid (TCA) precipitation. Greater than 95% of the total radioactivity counts were recovered upon TCA precipitation. The specific activity of ^125^I-Aβ40 and ^125^I-Aβ42 was determined to be in the range of 45–48 mCi/mg. The ^18^F-FDG was prepared in-house at the Mayo Clinic at a specific activity of 2-5 Ci/µmole as per the clinical standards.

### Kinetics of ^125^I-Aβ and ^18^F-FDG uptake in HFD versus LFD mice

#### Plasma pharmacokinetics and brain accumulation of ^125^I-Aβ and ^18^F-FDG

These experiments were carried out as described in our previous publications ^38–40^. Briefly, mice were anaesthetized and both femoral vein and artery were catheterized. A single intravenous bolus injection of either ^125^I-Aβ40 or ^125^I-Aβ42 (100 µCi/100 µL), or ^18^F-FDG (500 µCi/200 µL), was administered through the femoral vein. Blood samples (20 µL) were collected from the femoral artery at 0.25, 1, 3, 5, 7, 10, 15, 20, 30, and 40 minutes post-injection. The plasma was separated and was subjected to TCA precipitation. The ^125^I radioactivity levels in both the precipitate and supernatant were measured using a gamma counter (Cobra II; PerkinElmer Life and Analytical Sciences, Boston, MA). The radioactivity in the precipitate is attributed to intact protein, while the radioactivity in the supernatant is attributed to degraded protein. To assess ^18^F radioactivity, the blood samples were centrifuged to separate red blood cells and plasma, and the radioactivity of samples was assessed using the gamma counter. The ^18^F-FDG plasma time-activity [*C_p_*(*t*)] profiles were corrected for the ^18^F physical decay ^41–45^. As for ^125^I-Aβ40 and ^125^I-Aβ42, we did not adjust for decay due to their considerably long half-life of approximately 60 days.

The ^125^I-Aβ40, ^125^I-Aβ42, and ^18^F-FDG plasma concentration versus time profiles was fitted to a two-compartment model using Phoenix^®^ WinNonlin 6.4 (Certara, St. Louis, MO). The secondary plasma pharmacokinetics parameters, including Cmax, area under the curve (AUC), and clearance (CL) were also predicted.

#### Evaluation of ^125^I-Aβ brain influx using dynamic SPECT/CT imaging

The dynamic Single Photon Emission Computed Tomography/ Computed Tomography (SPECT/CT) imaging studies were performed as described in our earlier publications ^39, 46–48^. Briefly, the mice fed with LFD or HFD (n=4-5 per group) were anesthetized and catheterized as described above. A single dose of ^125^I-Aβ40 or ^125^I-Aβ42 (500 µCi in PBS) was administered by intravenous bolus injection via the femoral vein. The mouse was immediately transferred to a SPECT/CT scanner (Gamma Medica, Northridge, CA), and SPECT scans were taken at 1 min intervals over the next 40 min. This was followed by a 5 min CT scan to map the anatomical regions of interest (ROI). At the end of the imaging experiment, the mouse was transcardially perfused with excess PBS to flush any remaining ^125^I activity from the vasculature, and a final SPECT scan was taken to measure the ^125^I-Aβ tissue accumulation.

#### Assessment of ^18^F-FDG brain influx using dynamic PET/CT imaging

Mice fed with LFD or HFD were anesthetized as aforementioned. A bolus of ^18^F-FDG (500 μCi/200 µL) was injected into the femoral vein, and the mouse was imaged in the PET/CT scanner (Siemens Inveon® Multi-Modality System, Knoxville, TN) from 0 to 30 min followed by a 5 min CT scan to locate ROI.

#### Prediction of the brain influx rate of ^125^I-Aβ40, ^125^I-Aβ42, and ^18^FDG using dynamic SPECT and PET imaging data

The plasma-to-brain influx rate of ^125^I-Aβ40, ^125^I-Aβ42, and ^18^F-FDG in mice on LFD and HFD was quantified using Gjedde-Patlak graphical analysis ^49, 50^. The Gjedde-Patlak graph was constructed by plotting the ratio of the brain isotope activity normalized with the plasma concentration at time t, 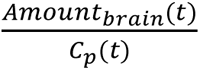, against the ratio of the cumulative area under the plasma concentration-time curve normalized to the plasma concentration at time, 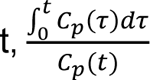. Here, *Amount_brain_*(*t*) denotes the isotope activity (µCi) within the brain ROI at time t, and *C_p_*(*t*) represents the plasma concentration (µCi/mL) at the same time point, whereas 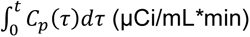 is the area under the plasma concentration versus time profile from 0 to t minutes, calculated using the trapezoidal method. The plasma-to-brain influx rate, K_i_ (mL/min) could be predicted from the slope of the linear regression of this plot.

#### The permeability-surface area (PS) products of ^125^I-Aβ40 and ^125^I-Aβ42 in the brains of LFD-versus HFD-fed mice

The PS products (mL/min/g) of ^125^I-Aβ uptake in the brain were calculated using the following equation:

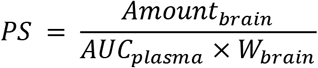

Where, *Amount_brain_* represents the total amount of ^125^I activity (µCi) in the brain regions post-perfusion, *AUC_plasma_* is the area under the plasma concentration-time curve (µCi·min/mL) from 0 to 40 min calculated using the trapezoidal method, and *W_brain_* denotes the weight of the brain region in grams (g).

#### Accumulation of ^18^F-F DG in the brain of LFD-versus HFD-fed mice

At the end of the ^18^F-FDG PET/CT study described above, the LFD and HFD mice were perfused with excess phosphate-buffered saline (PBS), then the brain was dissected and assayed for ^18^F activity using the gamma counter. The accumulation of ^18^F-FDG in the brain was expressed using the following equation:

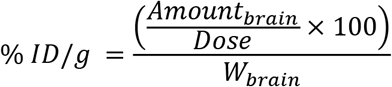

In this equation, % *ID*/*g* denotes the percentage of the injected dose per gram of brain tissue. Here, *Amount_brain_* is the amount of ^18^F activity (µCi) in the brain regions post-perfusion, *Dose* is approximately 500 µCi, and *W_brain_* is the brain weight (g).

### Cell culture

The human cerebral microvascular endothelial cell (hCMEC/D3) monolayers were used as a BBB model. These cell lines were a kind gift from P-O Couraud and were cultured as previously described ^51^.

### Brain microcapillaries isolation

As previously detailed by Gali CC. et al ^52^, brains were harvested from mice fed with either LFD or HFD. Briefly, 3 to 4 brain hemispheres were selected after removing meninges and olfactory bulbs. Then the brain hemispheres were homogenized in MCDB131 medium, which is supplemented with 2% FBS, 1% L-glutamine and 1% penicillin/streptomycin (Life Technologies, Carlsbad, CA). This brain homogenate was then digested for 1 hour at 37 °C using dispase (0.01 g per 2 hemispheres; Life Technologies), and subsequently centrifuged at 6,800 g on a dextran gradient using a 1:1 v/v dextran solution (1.0612 g/L; Alfa Aesar, Haverhill, MA). Then the cell suspension underwent an additional 1 minute of digestion at 37 °C with collagenase/dispase (3.5 mg/20 ml; Roche, Basel, Switzerland) to separate the basement membrane from the endothelium. The mixture was then filtered through 180 μm nylon mesh and centrifuged on a Percoll (Sigma-Aldrich, St. Louis, MO) density gradient to isolate pure microcapillary fragments from the interface. The final microvascular fragments were rinsed and stored at −80 °C for further analysis.

### Western blot

The brain microcapillaries or hCMEC/D3 cells were washed and lysed in RIPA buffer containing protease and phosphatase inhibitors. Lysates were loaded onto 4-12 % Criterion XT precast gels for separation by SDS-PAGE. After electrophoresis, proteins were transferred to a 0.2 µm nitrocellulose membrane. The membrane was then blocked and incubated with various primary antibodies (1: 500 for TXNIP and 1:000 for other proteins) overnight at 4 °C. On the next day, the membrane was washed and incubated with a fluorescent-conjugated secondary antibody (1:2000) for 1.25 hours at room temperature. The bands were imaged by LI-COR imaging system, and band intensity was quantified by Image Studio software.

### Cellular uptake of ^125^I-Aβ

To investigate the effects of insulin resistance on cellular uptake of Aβ peptides, the hCMEC/D3 cell monolayers were cultured on 12-well plates and pre-treated with either MEK inhibitor trametinib (1 µM) or AKT inhibitor MK2206 (12.5 µM) for 10 minutes, followed by a 20-minute stimulation with Humulin^®^. Subsequently, ^125^I-Aβ40 or ^125^I-Aβ42 were spiked into the medium (5 µCi/mL), and the cells were incubated for 60 minutes at 37 °C. After the treatment, the cells were washed and lysed using RIPA buffer. The radioactivity in the cell lysates was then measured using the gamma counter, as previously described.

### Fluorescent 2-NBDG cell uptake

To investigate the effects of insulin resistance on cellular uptake of glucose, the hCMEC/D3 cell monolayers were cultured on 6-well plates and pre-treated with either MEK inhibitor trametinib or AKT inhibitor MK2206, followed by stimulation with Humulin^®^. After the treatment, the cells were washed with ice-cold PBS, trypsinized, then centrifuged and resuspended in 440 µM of fluorescent D-glucose analog 2-[N-(7-nitrobenz-2-oxa-1,3-diazol-4-yl)amino]-2-deoxy-d-glucose (2-NBDG) in glucose-free cell medium for 30 minutes at 37 °C. Subsequently, the cells were centrifuged again, and washed twice with PBS, resuspended in PBS, and the median fluorescent intensity was measured by flow cytometry (BD Biosciences, San Jose, CA).

### Statistical analysis

Statistical tests were performed using GraphPad Prism 8.4 (GraphPad software; La Jolla, CA). For comparisons between HFD-fed and LFD-fed mice, a two-tailed student’s t-test was used.

## RESULTS

### Higher body weight and blood glucose in HFD-fed mice compared to LFD-fed mice

At 7-11 months of age, the mice were transitioned to either LFD or HFD, and their body weights were measured weekly. During 13-16 weeks on the LFD, there was an insignificant increase in the mouse body weight, from an initial mean weight of 33.5 ± 6.2 g to 34.2 ± 3.2 g. However, the mice on HFD showed a more pronounced increase in body weight, increasing to 1.6 times their original weight prior to the dietary modification (***Figure 1B***, p<0.001, two-tailed t-test). As expected, dietary modification has no impact on the brain weight (***Figure 1C***).

**Figure 1.**
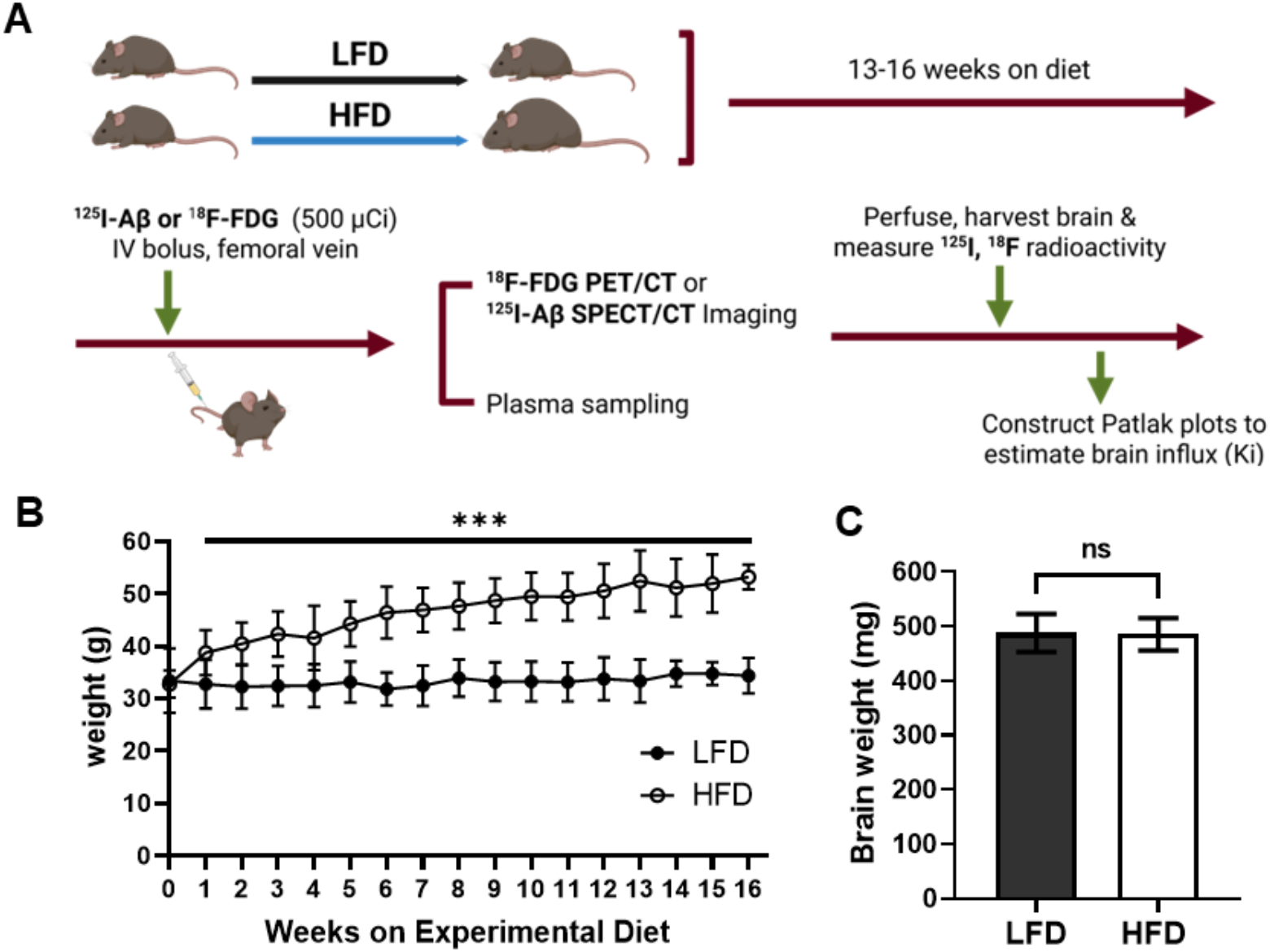
A high-fat diet (HFD) induces weight gain in B6SJLF1 mice compared to a low-fat diet (LFD). **(A)** Experimental design. B6SJLF1 mice were randomly allocated to 60% high-fat diet (HFD) or 10% low-fat diet (LFD) at 7-11 months of age for a period of 13-16 weeks. Those mice were then used for either dynamic SPECT/CT or PET/CT imaging or to assess plasma kinetics and brain permeability of ^125^I-Aβ or ^18^F-FDG under different diet conditions. Changes in **(B)** total body weight **(C)** brain weight after 13-16 weeks of HFD versus LFD. Mice on HFD showed an expected increase in body weight, but brain weight remained unchanged. Data are presented as mean ± SD (n=15 for LFD; n=15 for HFD); unpaired two-tailed Student’s t-test (**p<0.001).

To evaluate peripheral insulin sensitivity, an ITT was performed on mice fed with either LFD or HFD. Following the subcutaneous administration of insulin, a comparative analysis of blood glucose levels revealed that HFD-fed mice exhibited elevated glucose concentrations compared to their LFD-fed counterparts at 15-, 25-, and 45-minutes following insulin injection (***Figure2A***). Additionally, the AUC was calculated to capture cumulative glucose levels over a 90-minute period. The AUC for HFD-fed mice was 1.2 times greater compared to that of LFD-fed mice (p<0.001, two-tailed t-test).

### Plasma pharmacokinetics of ^125^I-Aβ are not substantially altered in HFD-fed mice compared to LFD-fed mice

After administering a single bolus dose of ^125^I-Aβ40 or ^125^I-Aβ42 into the femoral vein of mice fed with either a LFD or a HFD, a bi-exponential decrease in plasma concentrations of ^125^I-Aβ peptides was observed over time (***Figure 3A&B***). This pattern aligns with the outcomes reported in previous studies conducted on wild-type mice ^47, 53^. The peak plasma concentration (Cmax) of ^125^I-Aβ40 in HFD-fed mice was observed to be half (p<0.001, two-tailed t-test), and steady-state volume of distribution (Vss) was three-fold higher than that observed in LFD-fed mice (p<0.05, two-tailed t-test). Additionally, the plasma CL and AUC of ^125^I-Aβ40 was not significantly affected by the dietary changes. Interestingly, Cmax, AUC, and Vss of ^125^I-Aβ42 were relatively lower, and the plasma CL was found to be elevated in HFD-fed mice relative to LFD-fed mice. However, these differences were not statistically significant, thereby suggesting that dietary differences did not substantially alter the overall plasma pharmacokinetics of ^125^I-Aβ42.

**Figure 2.**
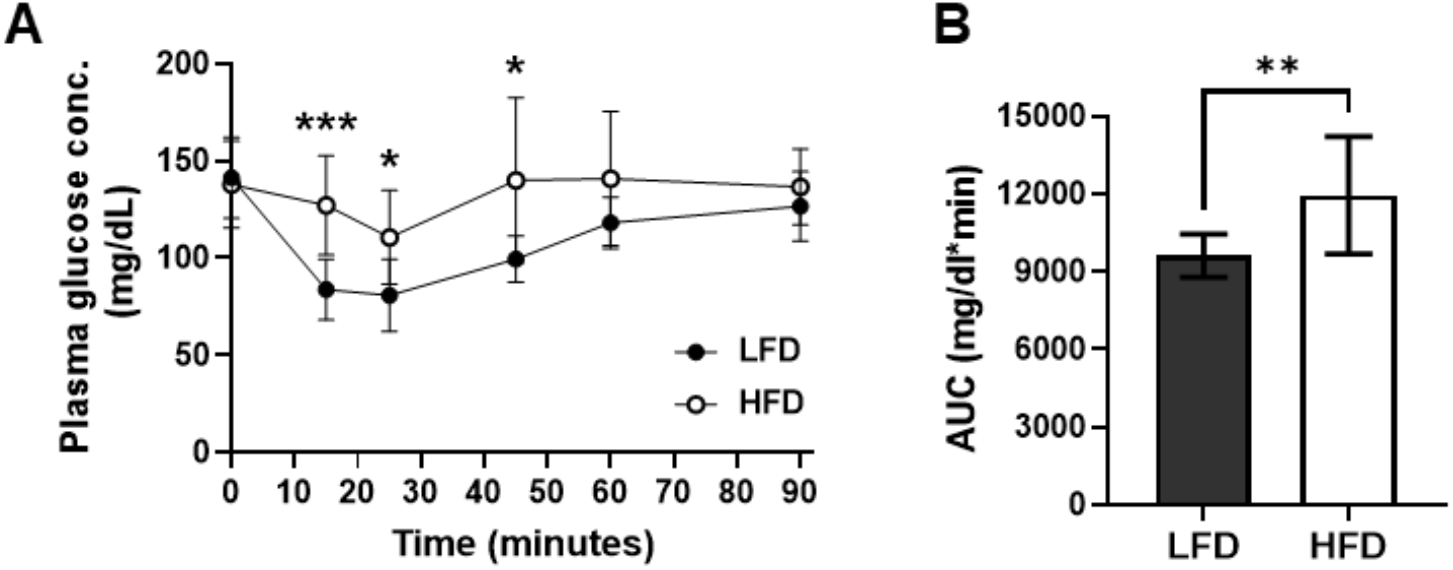
Insulin tolerance tests (ITTs) in high-fat diet (HFD)-fed and low-fat diet (LFD)-fed B6SJLF1 mice. **(A)** Changes in plasma glucose concentration over time following insulin injection. Data are represented as mean ± SD (n= 7-10). **(B)** After 13-16 weeks of HFD feeding, the mice demonstrated a higher ITT area under the curve (AUC) from 0-90 minutes, calculated from the plasma glucose concentration versus time profile, compared to LFD-fed mice. Data are represented as mean ± SD (n= 7-10); unpaired two-tailed Student’s t-test (**p<0.01).

**Figure 3.**
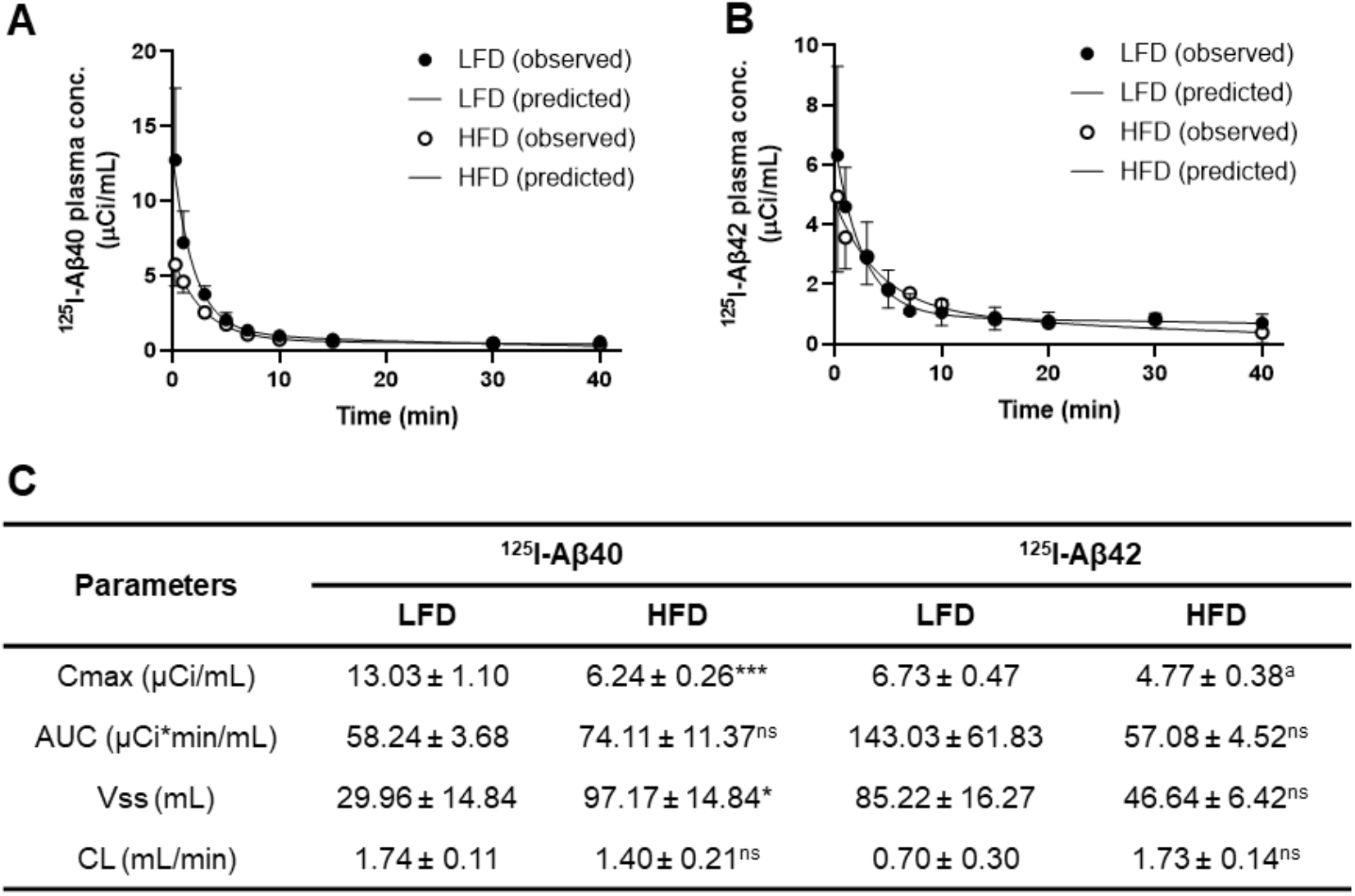
Plasma pharmacokinetics of ^125^I-Aβ peptides in high fat diet (HFD)-fed and low fat diet (LFD)-fed B6SJLF1 mice. HFD or LFD-fed mice were bolus injected with 100 µCi of ^125^I-Aβ into the femoral vein. Plasma was sampled between 0 and 40 minutes periodically via the femoral artery, and radioactivity of the intact ^125^I-Aβ was measured after trichloroacetic acid (TCA) precipitation in the gamma counter. The plasma concentration of **(A)** ^125^I-Aβ40 and **(B)** ^125^I-Aβ42 vs. time data was fit to a bi-exponential equation. Data are represented as the observed values (mean ± SD; ^125^I-Aβ40: n=6 for LFD and n=5 for HFD; ^125^I-Aβ42: n=13 for LFD and n= 3 for HFD) superimposed on the predicted curves. **(C)** Pharmacokinetic parameters predicted for ^125^I-Aβ40 (n=6 for LFD and n=5 for HFD) and ^125^I-Aβ42 (n=13 for LFD and n= 3 for HFD) in HFD-fed and LFD-fed B6SJLF1 mice. Data are represented as mean ± SEM; unpaired two-tailed Student’s t-test (*p<0.05, ***p<0.001, ^a^p=0.075; ns: not significant).

### Higher plasma-to-brain uptake of ^125^I-Aβ40 and ^125^I-Aβ42 in HFD-fed mice than in LFD-fed mice

The plasma-to-brain influx rate (Ki) of both ^125^I-Aβ40 and ^125^I-Aβ42, as determined by Patlak plot analysis, was higher in HFD-fed mice than in LFD-fed mice. For ^125^I-Aβ40, the Ki was 34.5 * 10^-4^ ± 4.9 * 10^-4^ mL/min in HFD-fed mice compared with 14.8 * 10^-4^ ± 3.3 * 10^-4^ mL/min in LFD-fed mice (***Figure 4A&C***). Similarly, the Ki for ^125^I-Aβ42 was 13.2 * 10^-4^ ± 3.3 * 10^-4^ mL/min in HFD-fed mice, which was significantly higher (p<0.001, two-tailed t-test) than that in LFD-fed mice (8.2 * 10^-4^ ± 1.7 * 10^-4^ mL/min) (***Figure 4B&C***). Consistent with this finding, the PS values for both ^125^I-Aβ40 (***Figure 5A***) and ^125^I-Aβ42 (***Figure 5B***) in HFD-fed mice were approximately twice as much as those found in LFD-fed mice (p<0.05, two-tailed t-test).

**Figure 4.**
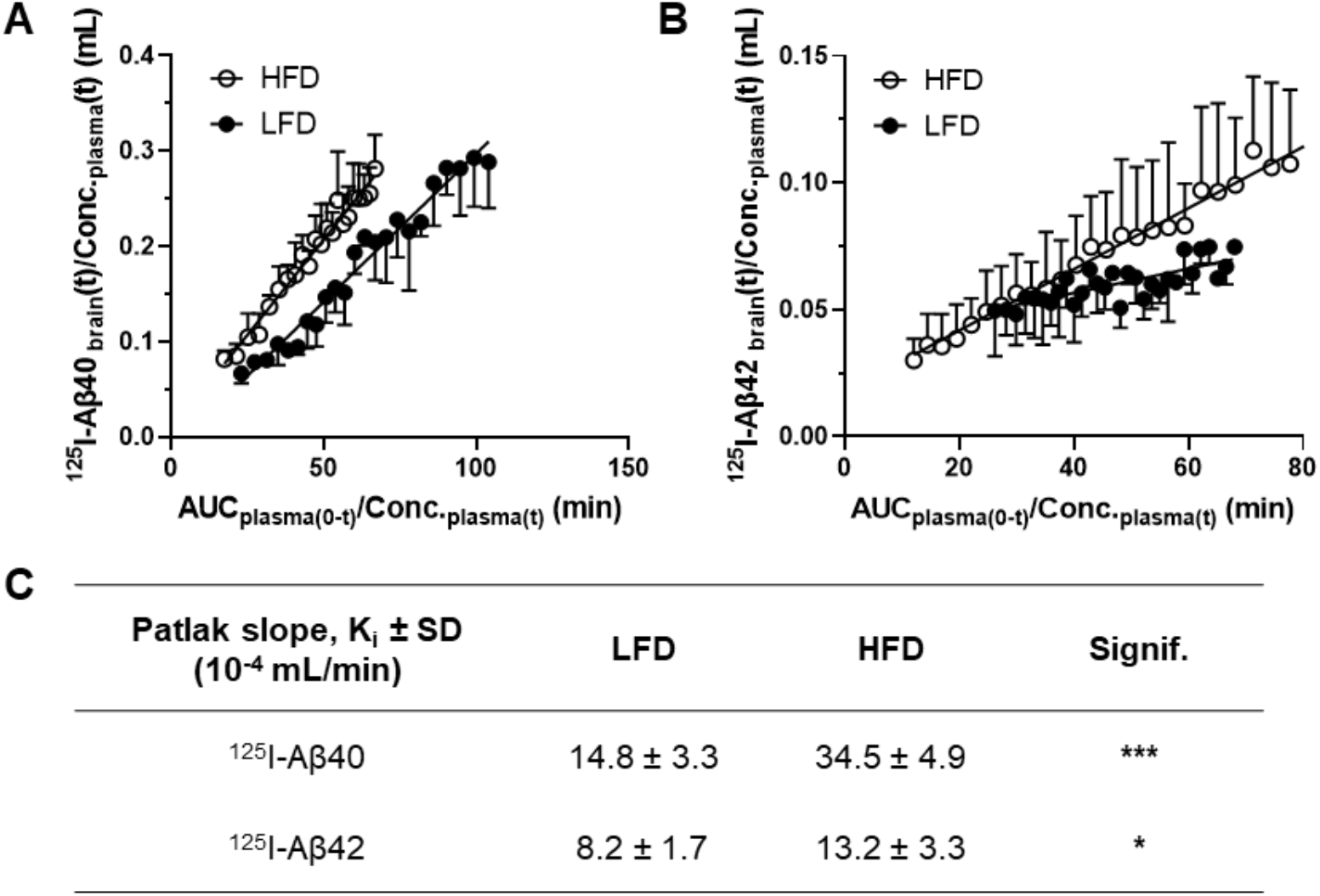
Impact of high-fat diet (HFD)-induced insulin resistance on brain Influx rate of ^125^I-Aβ40 and ^125^I-Aβ42 in Mice. Mice fed with LFD and HFD were administered with ^125^I-Aβ peptides (500 µCi) via the femoral vein, and radioactivity in the brain was monitored by dynamic SPECT/CT imaging from 0 to 40 min. Gjedde-Patlak plots show brain influx rate of **(A)** ^125^I-Aβ40 and **(B)** ^125^I-Aβ42 in the brain of HFD-fed versus LFD-fed mice. **(C)** The brain influx rate (Ki) of ^125^I-Aβ40 and ^125^I-Aβ42 was estimated by the slope obtained from Gjedde-Patlak graphical analysis. Data are presented as mean ± SD (^125^I-Aβ40: n=4 for LFD and n=8 for HFD; ^125^I-Aβ42: n=4 for LFD and n=8 for HFD); unpaired two-tailed Student’s t-test (*p<0.05, ***p<0.001).

**Figure 5.**
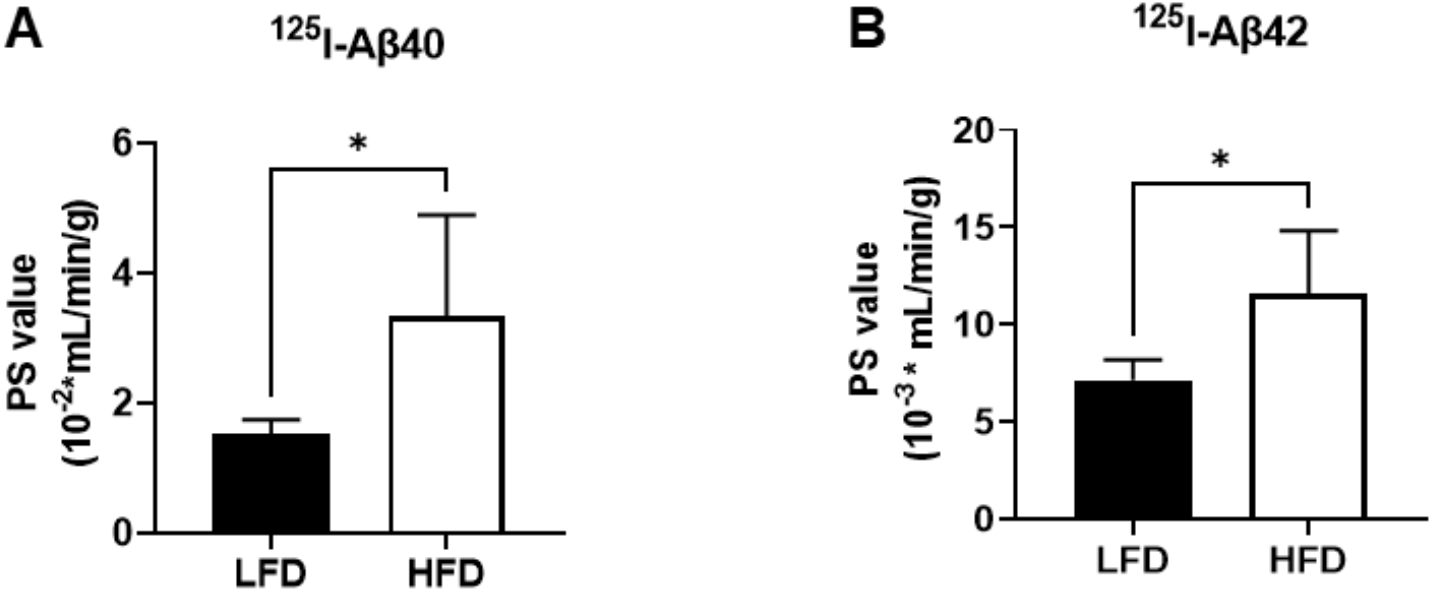
Brain permeability of ^125^I-Aβ40 and ^125^I-Aβ42 is increased in HFD-fed mice compared to LFD-fed mice. The permeability-surface area (PS) products for **(A)** ^125^I-Aβ40 and **(B)** ^125^I-Aβ42 uptake at the BBB in HFD-fed mice versus LFD-fed mice. Data are presented as mean ± SD (^125^I-Aβ40: n=4 for LFD and n=8 for HFD; ^125^I-Aβ42: n=4 for LFD and n=8 for HFD); unpaired two-tailed student t-test (*p<0.05).

### Lower plasma-to-brain uptake of ^18^F-FDG without affecting its plasma disposition in HFD-fed mice compared to LFD-fed mice

The plasma concentration of ^18^F-FDG exhibits a bi-exponential decrease over time (***Figure 6A&B***) and the observed concentrations in LFD- and HFD-fed mice were not significantly different. However, the HFD-fed mice demonstrated lower plasma-to-brain influx rate (18.3 * 10^-3^ ± 3.2 * 10^-3^ mL/min) compared to LFD-fed mice (8.7 * 10^-3^ ± 1.6 * 10^-3^ mL/min) (***Figure 7A***, ***p<0.01, two-tailed t-test). The accumulation of ^18^F-FDG in brain upon intravenous administration was determined as the percentage of the injected dose per gram of brain tissue (% ID/g), and was found to be lower in HFD-fed mice compared to LFD-fed mice (***Figure 7B***, p<0.05, two-tailed t-test).

**Figure 6.**
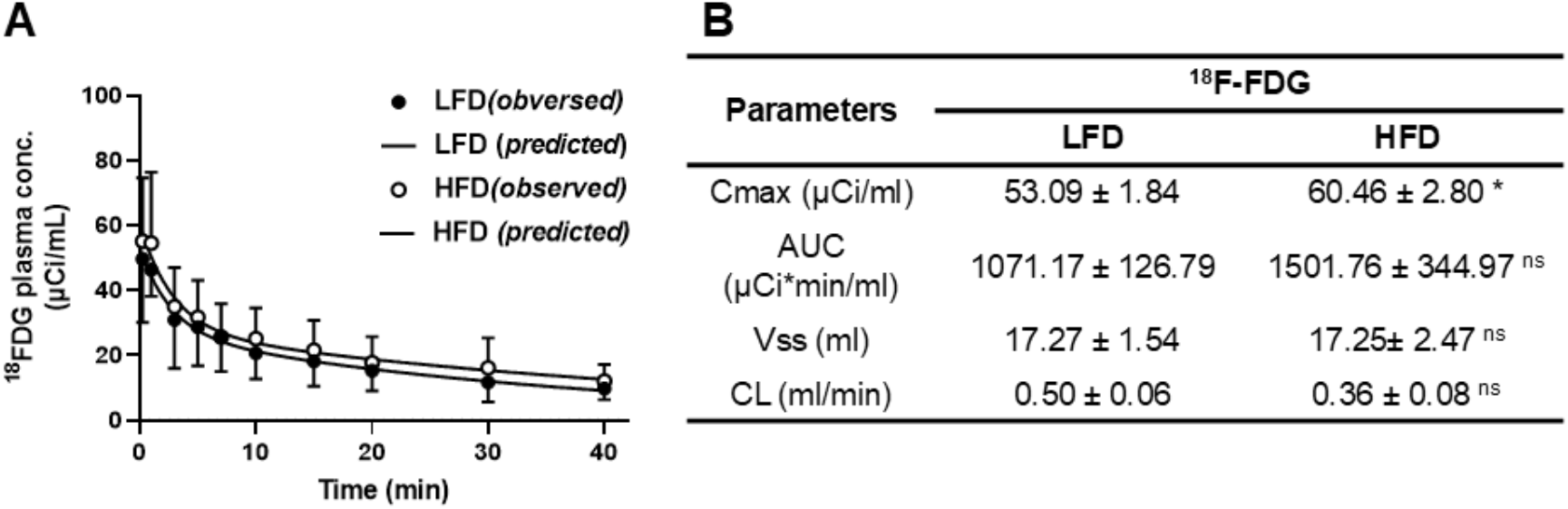
Plasma pharmacokinetics of ^18^F-FDG in HFD-fed and LFD-fed B6SJLF1 mice. HFD or LFD fed mice were bolus injected with 500 µCi of ^18^F-FDG via the femoral vein. Plasma was sampled periodically between 0-40 minutes, and radioactivity was measured by gamma counter. **(A)** The plasma concentration of ^18^F-FDG versus time data was fit to a bi-exponential equation. Data are presented as the observed values (mean ± SD) for both LFD (n=7) and HFD (n=5) overlaid on the predicted curves. **(B)** Pharmacokinetic parameters of ^18^F-FDG in HFD-fed and LFD-fed B6SJLF1 mice. Data are presented as mean ± SEM (n=7 for LFD and n=5 for HFD); unpaired two-tailed student’s t-test (*p<0.05, ns: not significant).

**Figure 7.**
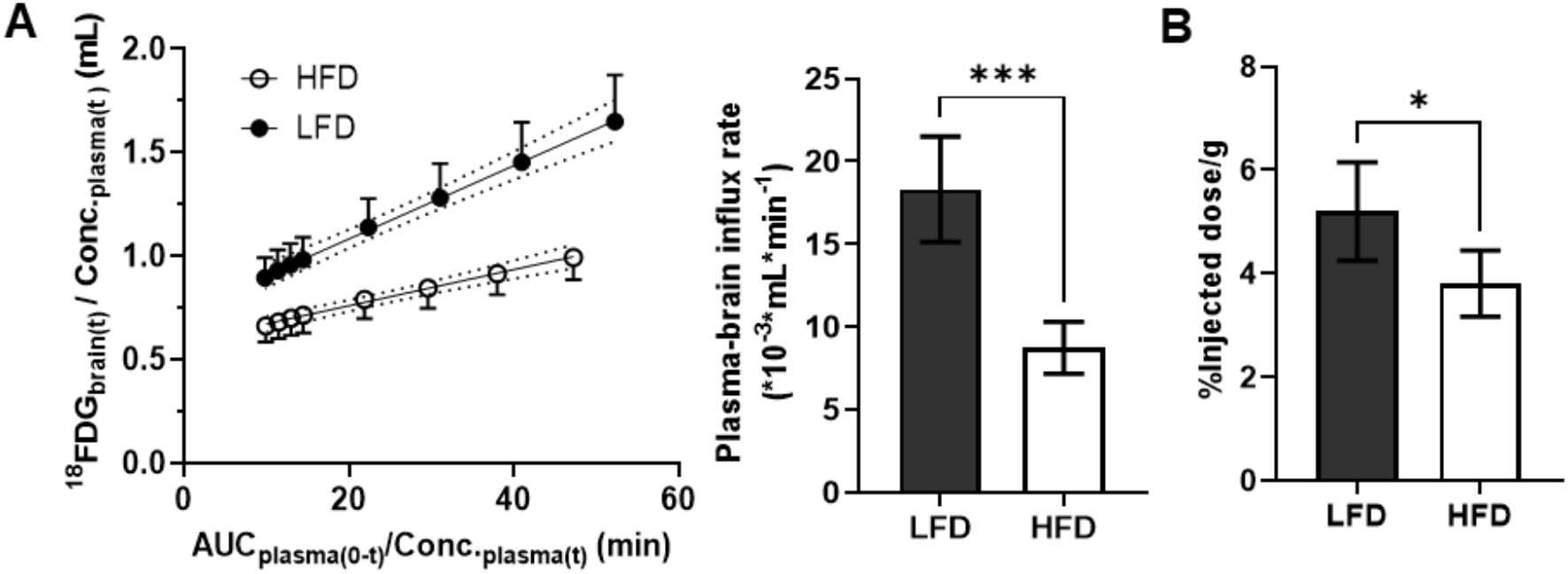
Impact of HFD-Induced Insulin Resistance on Brain Influx rate of ^18^F-FDG in Mice. Mice fed with LFD and HFD were administered ^18^F-FDG peptides (500 µCi) via femoral vein and its brain accumulation was monitored by dynamic PET/CT imaging from 0-30 min. **(A)** The brain influx rate (Ki) of ^18^F-FDG was estimated by the slope obtained from Gjedde-Patlak graphical analysis in HFD-fed and LFD-fed mice. Data are presented as mean ± SD (n=5 for both LFD and HFD); paired two-tailed student’s t-test (***p<0.01). **(B)** After PET/CT imaging, the brain accumulation of ^18^F-FDG in LFD-fed and HFD-fed mice was measured by gamma counter and presented as the percentage of the injected dose accumulating in the brain. Data are presented as mean ± SD (n=8 for both LFD and HFD); paired two-tailed student’s t-test (*p<0.05).

### Disruption of transporter expression and insulin signaling in the brain microcapillaries of HFD-fed mice compared to LFD-fed mice

Given the previously reported roles of RAGE in facilitating the plasma-to-brain entry of Aβ and GLUT1 in mediating glucose transport across the blood-brain barrier (BBB), we investigated if RAGE and GLUT1 expressions are altered in brain microcapillaries obtained from HFD-fed mice compared to LFD-fed mice. We found an upregulation of RAGE expression and downregulation of GLUT1 in mice fed with HFD (***Figure 8A-C***, p<0.05, two-tailed t-test). Moreover, we found no significant difference in the expressions of putative Aβ transporters, such as LRP-1 and P-glycoprotein (P-gp), in the brain microcapillaries obtained from LFD-fed versus HFD-fed mice.

**Figure 8.**
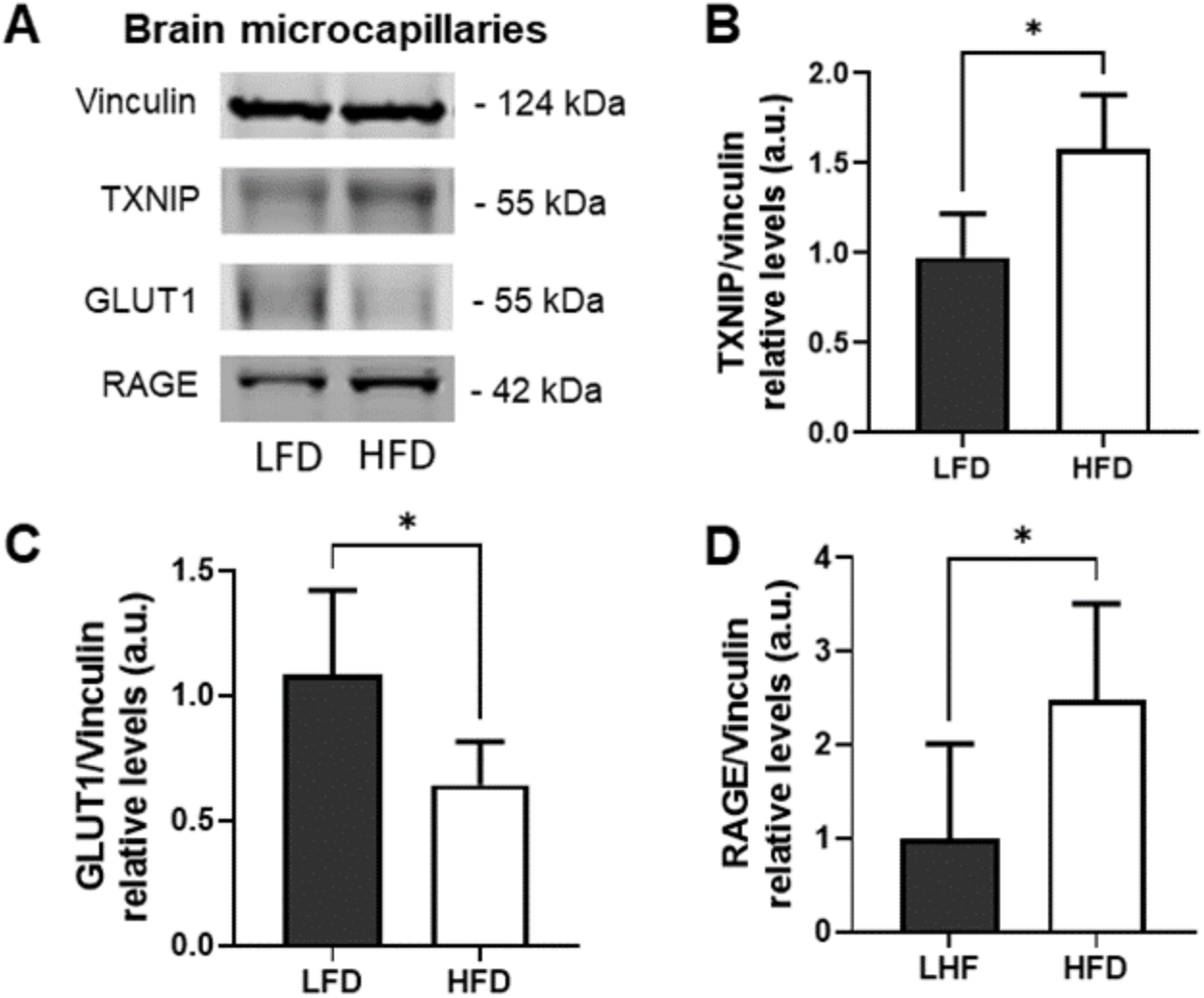
Western blots for Aβ influx transporter RAGE, glucose transporter (GLUT1), and negative regulator of GLUT1 expression, TXNIP, in brain microcapillaries harvested from LFD-fed and HFD-fed mice. **(A)** Representative blots are shown. Bar charts representing the quantification of expressions of **(B)** TXNIP/Vinculin, **(C)** GLUT1/Vinculin, and **(D)** RAGE/Vinculin. Data are presented as Mean ± SD (n=5 for LFD and n=4 for HFD); unpaired two-tailed student’s t-test (*p<0.05).

Our previous research has identified TXNIP as a negative regulator of GLUT1 ^40^. Hence, we assessed the TXNIP expression in mouse brain microvessels and observed an increase in TXNIP expression in the brain microvessels obtained from HFD-fed mice compared to their LFD-fed counterparts (***Figure 8A&D***, p<0.05, two-tailed t-test). Moreover, microvessels from HFD-fed mice demonstrated a decrease in AKT and ERK phosphorylation compared to those obtained from LFD-fed mice (***Figure 9***, p<0.01, two-tailed t-test).

**Figure 9.**
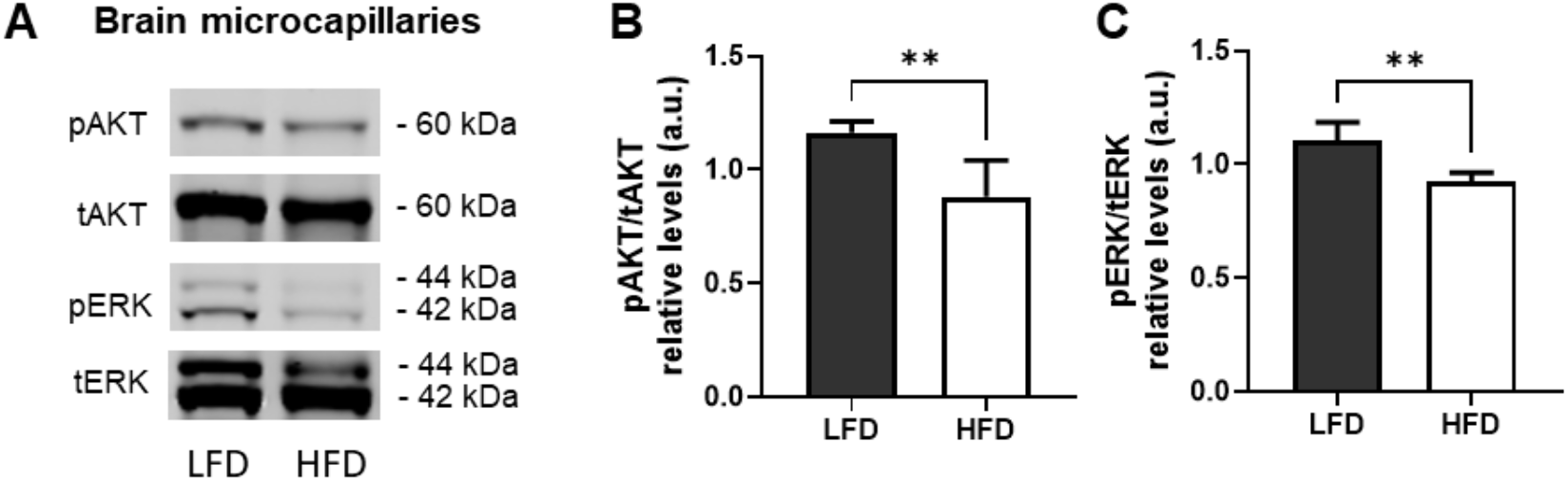
Expression of phospho-AKT (pAKT) and phospho-ERK (pERK) in brain microcapillaries harvested from LFD-fed and HFD-fed mice. **(A)** Representative blots are shown. Bar chart showing the quantification for **(B)** phospho-AKT(pAKT)/total AKT (tAKT) and **(C)** phosphor-ERK (pERK)/total ERK (tERK). Data are presented Mean ± SD (n=5 for LFD and n=4 for HFD); unpaired two-tailed student’s t-test (**p<0.01).

### MEK and AKT inhibitors differentially effect ^125^I-Aβ and fluorescent glucose analog (2-NBDG) uptake in hCMEC/D3 cell monolayers

The MEK inhibitor trametinib and AKT inhibitor MK2206 have been shown to specifically reduce the phosphorylation of ERK and AKT, respectively (***Figure 10***). Treatment with trametinib led to a 1.2-fold increase in the cellular uptake of ^125^I-Aβ40 (***Figure 11A***, p<0.05, two-tailed t-test), but demonstrated no significant effect on the uptake of ^125^I-Aβ42 (***Figure 11C***). Conversely, MK2206 treatment resulted in an increase in the cellular uptake of ^125^I-Aβ42 (***Figure 11D***, p<0.05, two-tailed t-test), but showed no significant change in the uptake of ^125^I-Aβ40 (***Figure 11B***). Furthermore, both trametinib (***Figure 11E&F***, p<0.0001, two-tailed t-test) and MK2206 (***Figure 11G&H***, p<0.05, two-tailed t-test) treatments were found to reduce the cellular uptake of 2-NBDG in hCMEC/D3 cell monolayers.

**Figure 10.**
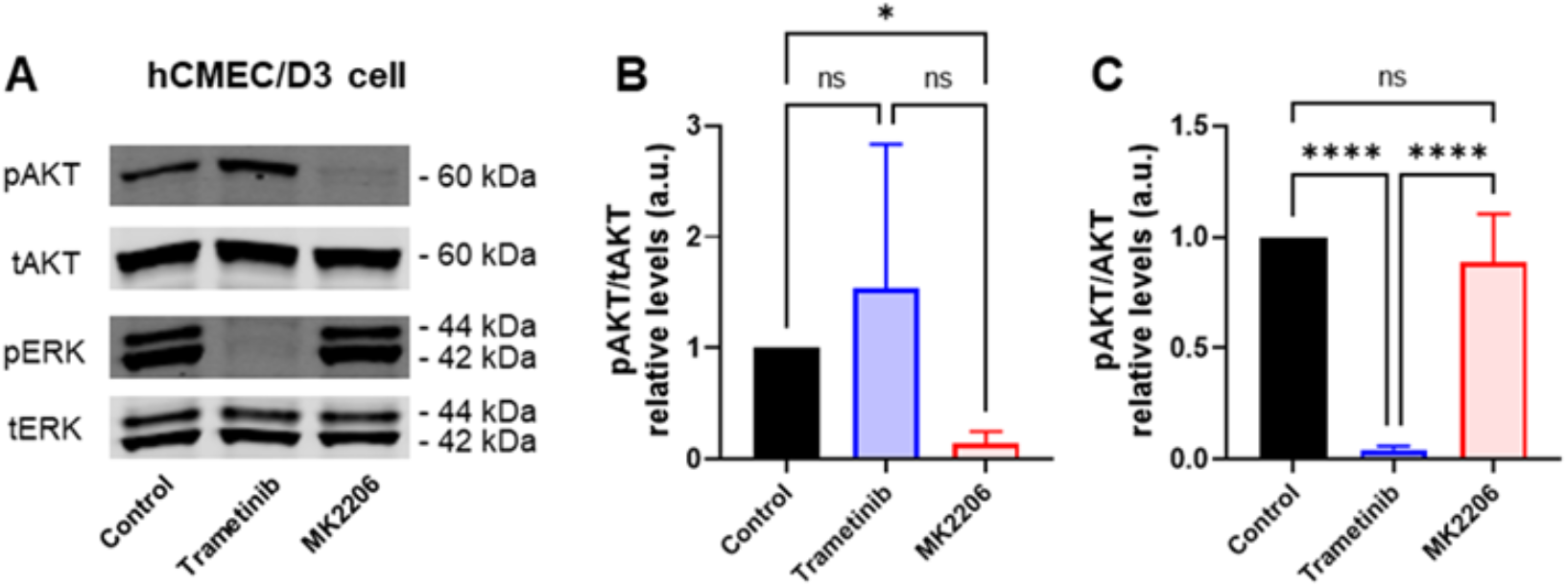
Effects of MEK and AKT inhibitors on the expression of pAKT and pERK in BBB endothelial cell monolayers. The hCMEC/D3 cell monolayers were pretreated with either 1 µM trametinib (MEK inhibitor) or 12.5 µM of MK2206 (AKT) inhibitor, for 10 minutes, followed by a 20-minute stimulation with 100 nM of insulin (Humulin^®^). Then the lysates were collected for western blots. **(A)** Representative blots showing the expressions of phosphor-AKT (pAKT), total AKT, phosph-ERK (pERK), and total ERK (tERK). Bar chart of the quantification of (**B)** pAKT expression normalized to tAKT and **(C)** pERK expression normalized to tERK. Data are presented mean ± SD (n=3); paired two-tailed student t-test (*p<0.05; ****p<0.0001).

**Figure 11.**
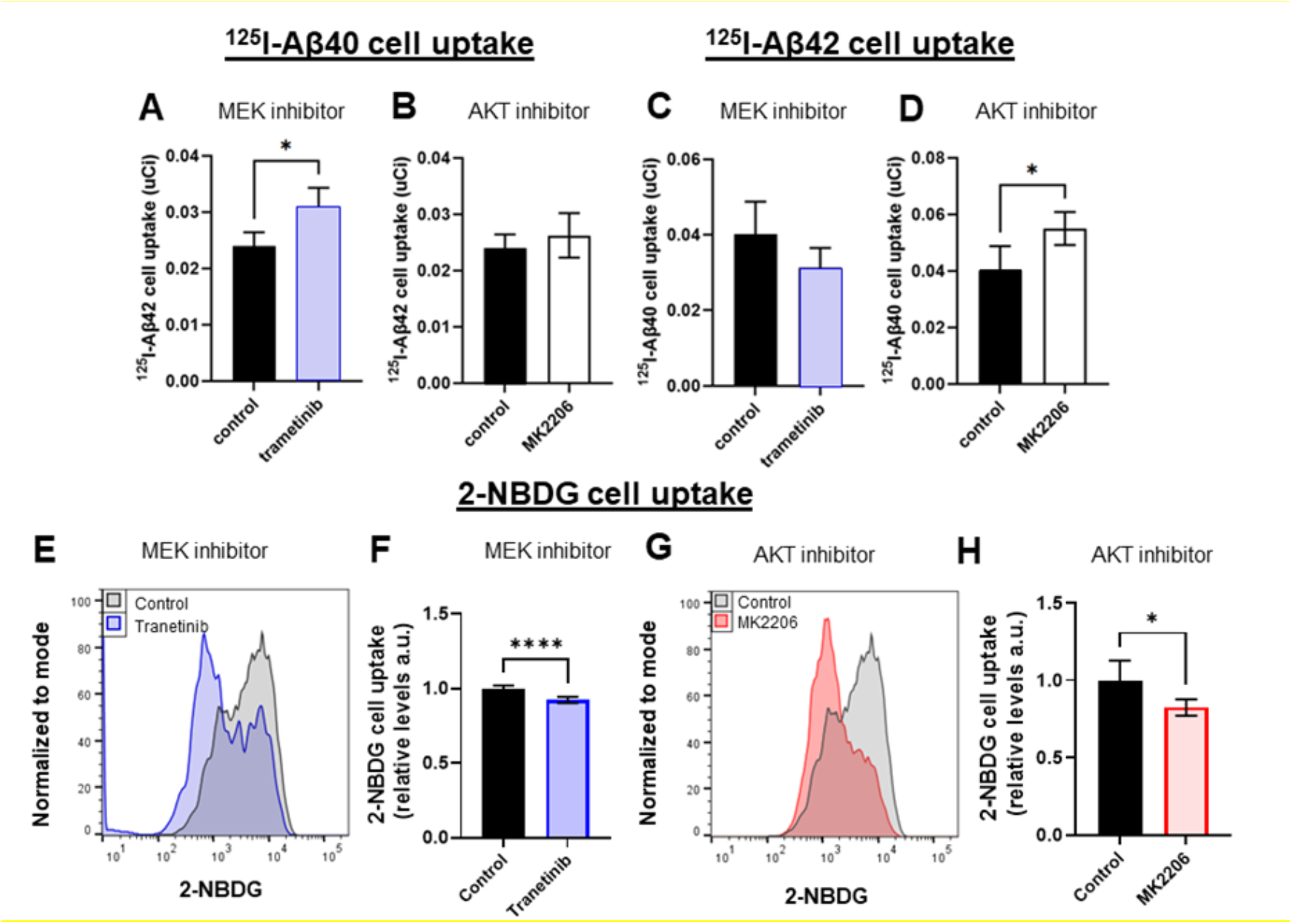
Effects of MEK and AKT inhibitors on Aβ40, Aβ42, and 2-NBDG (fluorescent glucose tracer) uptake in BBB endothelial cell monolayers. Cellular uptake of **(A&B)** ^125^I-Aβ40, **(C&D)** ^125^I-Aβ42, and **(E&H)** 2-NBDG were assessed in hCMEC/D3 cell monolayers upon insulin stimulation following treatment with various insulin signaling inhibitors. Specifically, cells were pretreated with either 1 µM MEK inhibitor trametinib or 12.5 µM AKT inhibitor MK2206, for 10 minutes, followed by a 20-minute stimulation with 100 nM insulin (Humulin®). Subsequently, cells were co-incubated with either ^125^I-Aβ (5 µCi/mL) or 2-NBDG (440 µM) for another 1 hour. The radioactivity of ^125^I-Aβ was measured by gamma counter, while the mean intensity signal of 2-NBDG was measured by flow cytometry. Data are presented as Mean ± SD (n=3 for ^125^I-Aβ and n=6 for 2-NBDG); unpaired two-tailed student t-test (*p<0.05; ****p<0.0001).

## Discussion

Although metabolic syndrome and T2DM are major risk factors for AD onset and progression, the underlying pathological mechanisms remain incompletely understood. Several studies have shown that these risk factors exacerbate vascular dysfunction, which coexists with AD pathology in most patients, and accelerates cognitive decline. Vascular dysfunction is characterized by increased BBB permeability due to altered expression of tight junction proteins, impaired vascular responses, and loss of selectivity in BBB transport ^54, 55^. We hypothesized that insulin resistance, induced by feeding HFD, may play a pivotal role in BBB dysfunction, and exacerbate pathophysiological changes associated with AD.

In this study, we found that 7 to 11 months-old mice demonstrated a significant increase in the body weight as early as one week after initiation of HFD feeding (***Figure1B***). More importantly, a reduction in peripheral insulin sensitivity, as determined by an insulin tolerance test, was also observed after 13-16 weeks of HFD feeding (***Figure2***), confirming the development of diet-induced metabolic syndrome.

Although previous studies conducted in AD transgenic mouse models have shown that HFD contributes to increased amyloid accumulation, these altered Aβ levels are not consistently observed and may vary depending on factors such as sex, mouse model, type of dietary modification, and duration of HFD exposure ^56^. To evaluate the impact of HFD-induced peripheral insulin resistance on brain Aβ uptake, we employed radiolabeled Aβ tracers to monitor Aβ trafficking at the BBB. Patlak plots describing the influx rate into the brain have indicated higher brain influx rate (***Figure5***) of both ^125^I-Aβ40 and ^125^I-Aβ42 and greater brain accumulation in HFD-fed mice compared to LFD-fed mice. However, the systemic exposure, as indicated by plasma AUC of ^125^I-Aβ40 or ^125^I-Aβ42 showed no significant difference between LFD-fed and HFD-fed mice (***Figure3***). This indicates that HFD-induced peripheral insulin resistance does not affect Aβ exposure to the BBB endothelium. Hence, we investigated changes in the expression of putative Aβ transporters at the BBB endothelium. Indeed, RAGE, which mediates Aβ influx, was upregulated in the brain microvessels of HFD-fed mice (***Figure8***), while no changes were observed in the expressions of LRP1 or P-gp, both of which facilitate Aβ efflux from the brain ^57^. Previous studies have shown elevated RAGE expression in diabetic models, although they primarily focused on type-1 diabetes characterized by irreversible damage to pancreatic islets ^58^ and did not specifically investigate RAGE at the BBB endothelium^59^.

Our focus is primarily on pathological changes associated with T2DM, which manifests hyperinsulinemia, hyperglycemia, and dyslipidemia. T2DM can be induced genetically or through dietary modification by supplementing it with higher fat content compared to the LFD. However, genetic mutations such as leptin deficiency that trigger metabolic syndrome are uncommon in AD patients ^60^. Nonetheless, the risk of metabolic syndrome may significantly increase in AD patients due to unhealthy lifestyle, including excessive caloric consumption or physical inactivity. Therefore, diet induced T2DM model is a better representation of the metabolic changes observed in AD patients. Additionally, previous studies have shown that an increase in Aβ production does not fully account for the increase in Aβ deposition observed in AD patients. Instead, a decrease in the clearance of Aβ from the brain parenchyma and/or increase of Aβ from in the systemic circulation may play a more significant role ^61–63^.

Consistent with our findings, previous studies have shown that RAGE expression increased in the cortex of HFD-fed mice ^5^, but its BBB-specific expression was not explored in this study. Moreover, studies have shown no significant changes in both protein and mRNA levels of P-gp in the brain microvessels of mice after placing them two months on HFD ^7^. Similarly, LRP1 expression remained unchanged in both WT and AD transgenic mice after three months on HFD ^64^. Overall, we have identified a potential correlation between diet-induced metabolic alternations and the dysregulation of Aβ influx from plasma to the brain in HFD-fed mice.

Previous studies have shown that brain insulin resistance is characterized by impaired insulin signaling in the BBB endothelium ^62^. In addition, it has been suggested that insulin signaling regulates Aβ transcytosis across the BBB ^65^. In AD transgenic (3xTg) mice fed with HFD, which induces peripheral insulin resistance, there was a notable elevation in the levels of soluble Aβ40 and Aβ42 in the cerebral cortex. Conversely, the administration of insulin via intravenous injection mitigated the elevated soluble Aβ levels in the cortext caused by the HFD, implicating deficits in insulin signaling triggered by HFD may contribute to increased Aβ uptake at the BBB ^65^. Thus, we hypothesized that insulin signaling disruption, specifically in PI3K/Akt and MEK/Erk pathways, is responsible for enhanced Aβ accumulation observed in HFD mice. To test the hypothesis, we investigated the effect of various insulin signaling inhibitors on Aβ uptake. We found that treatment with MEK inhibitor trametinib increased endothelial uptake of ^125^I-Aβ42, whereas the uptake of ^125^I-Aβ40 remained unaffected (***Figure11A&C***). Strikingly, treatment with the AKT inhibitor, MK2206, did not alter the endothelial uptake of ^125^I-Aβ42 but increased the uptake of ^125^I-Aβ40 (***Figure11B&D***). Previous research from our lab also demonstrated that treating healthy mice with AG1024, which is an IGF1/IR dual kinase inhibitor ^66^ of both PI3K/Akt and MEK/Erk pathways, reduced influx rate as well as uptake of ^125^I-Aβ42 but did not impact ^125^I-Aβ40 influx into the brain ^67^. These observations support the hypothesis that impairments in insulin signaling are responsible for selectively disrupting Aβ40 versus Aβ42 trafficking at the BBB and contribute to greater Aβ deposition observed in HFD-fed mice.

T2DM is also associated with diminished cerebral glucose metabolism, especially in brain regions such as posterior cingulate cortex, the precuneus region, parietal cortices, in which a lower cerebral metabolic rate of glucose (CMRglc) was associated with greater insulin resistance ^27^. In our study, the mice that were fed with HFD for 4-months have also demonstrated reduced influx rate of ^18^F-FDG compared to those on LFD (***Figure7***).

Our analysis of brain microcapillaries obtained from HFD-fed mice has demonstrated a reduction in GLUT1 expression compared to microcapillaries obtained from LFD-fed mice (***Figure8***). This suggests that HFD may lead to diminished glucose uptake by reducing GLUT1 expression at the BBB. Previously, we identified TXNIP protein as a negative regulator of GLUT1 expression^40^. Consistent with this finding, we also noted an increase in TXNIP expression in the brain microcapillaries of HFD-fed mice compared to those obtained from LFD-fed mice (***Figure8***). These findings suggest that HFD could potentially reduce glucose uptake by reducing the expression of GLUT1 at the BBB. However, the reduced glucose influx rate could also be due to the reduction in cerebral perfusion, as documented in patients with insulin resistance and T2DM ^68–72^. Research investigating the changes in cerebral perfusion due to HFD is limited and the contribution of cerebral perfusion changes to glucose transport at the BBB warrants further investigation.

We further investigated the effect of insulin signaling disruption caused by HFD within brain microvessels on glucose transport to the brain. Unlike in peripheral tissues, glucose transport at the BBB is insulin-independent ^73^. However, insulin signaling pathways may influence GLUT1 distribution at the BBB through the regulation of TXNIP^74^, and indirectly affect plasma-to-brain glucose transport. This hypothesis is tested by investigating the effects of insulin signaling inhibitors on glucose uptake. As a result, we observed that the uptake of 2NBDG, a glucose surrogate, was diminished in the polarized BBB endothelial monolayers upon MEK or AKT inhibition (***Figure11***). These results imply that brain insulin resistance, triggered by HFD, may lead to diminished glucose uptake into the brain.

The AD pathology is aggravated by comorbidities such as diabetes and metabolic syndrome that are prevalent in the elderly^75^. The findings of the current study mechanistically connect two of the most visible changes in the AD brain captured by diagnostic imaging, Aβ deposition and hypometabolism, with insulin resistance observed in T2DM and metabolic syndrome. Further studies are needed to establish pathological synergism between Aβ deposition and insulin resistance in aggravating BBB dysfunction and worsening neuropathological changes in AD brain.

## List of abbreviations

LOAD: Late-onset Alzheimer’s disease
AD: Alzheimer’s disease
HFD: High-fat diet
Aβ: Amyloid-beta
T2DM: Type-2 diabetes mellitus
BBB: Blood-brain barrier
RAGE: Receptor for advanced glycation end products
LRP1: Lipoprotein receptor-related protein 1
P-gp: P-glycoprotein
ABC: ATP-biding cassette
CMRglc: Cerebral metabolic rate of glucose
GLUT1: Glucose transporter 1
Aβ: Amyloid-beta
^18^F-FDG: 18fluoredeoxyglucose
LFD: Low-fat diet
ITT: Insulin tolerance test
TCA: Trichloroacetic acid
AUC: Area under the curve
CL: Clearance
SPECT/CT: Single Photon Emission Computed Tomography/ Computed Tomography
PS: Permeability-surface area
PBS: Phosphate-buffered saline
hCMEC/D3: Human cerebral microvascular endothelial cell
2-NBDG: 2-[N-(7-nitrobenz-2-oxa-1,3-diazol-4-yl) amino]-2-deoxy-d-glucose
Cmax: Peak plasma concentration
Vss: Steady-state volume of distribution

## Author Contributions

Lushan Wang (Conceptualization; methodology; Formal Analysis; Investigation; Writing – Original Draft; Writing – Review & Editing); Geoffry L. Curran (Methodology; Investigation); Chaitanya Chakravarthi Gali (Methodology; Investigation); Andrew L. Zhou (Methodology; Formal analysis; Investigation); Paul H. Min (Conceptualization; Supervision; Project administration); Val J. Lowe (Conceptualization; Supervision; Project administration); Karunya K. Kandimalla (Conceptualization; Writing – Review & Editing; Supervision; Project administration; Funding acquisition)

## Acknowledgment

Figure 1A was generated using BioRender.

